# Evidence of signaling and adhesion roles for β-catenin in the sponge Ephydatia muelleri

**DOI:** 10.1101/164012

**Authors:** Klaske J. Schippers, Scott A. Nichols

**Author notes:** Correspondence: Scott. A. Nichols.

## Abstract

β-catenin acts as a transcriptional co-activator in the Wnt/β-catenin signaling pathway and a cytoplasmic effector in cadherin-based cell adhesion. These functions are ancient within animals, but the earliest steps in β-catenin evolution remain unresolved due to limited data from key lineages – sponges, ctenophores and placozoans. Previous studies in sponges have characterized β-catenin expression dynamics and used GSK3B antagonists to ectopically activate the Wnt/β-catenin pathway; both approaches rely upon untested assumptions about the conservation of β-catenin function and regulation in sponges. Here, we test these assumptions using an antibody raised against β-catenin from the sponge *Ephydatia muelleri*. We find that cadherin-complex genes co-precipitate with endogenous *Em* β-catenin from cell lysates, but that Wnt pathway components do not. However, through immunostaining we detect both cell boundary and nuclear populations, and we find evidence that *Em* β-catenin is a conserved substrate of GSK3B. Collectively, these data support conserved roles for *Em* β-catenin in both cell adhesion and Wnt signaling. Additionally, we find evidence for an *Em* β-catenin population associated with the distal ends of F-actin stress fibers in apparent cell-substrate adhesion structures that resemble focal adhesions. This finding suggests a fundamental difference in the adhesion properties of sponge tissues relative to other animals, in which the adhesion functions of β-catenin are typically restricted to cell-cell adhesions.

## INTRODUCTION

A well-known pattern in the fossil record is the appearance of most animal lineages over a relatively short time interval (between 541-485 Mya) during the Cambrian. Many of the characteristics of modern animals had already evolved by the time that these first body fossils appear, leaving few clues about either the process or sequence of animal body plan diversification. As a result, our understanding of the earliest events in animal evolution rests largely upon the comparative study of extant organisms. The rationale for this approach is that body plan differences between living organisms reflect accumulated evolutionary changes to their underlying cell and developmental biology.

For practical reasons, large research communities have established around a few genetically manipulable research models such as mouse, fruit fly, roundworm and zebrafish. Data from these organisms largely form the basis for textbook generalizations about animal biology, and have contributed to a bilaterian-biased perspective on animal evolution. However, to study the deepest periods of animal ancestry it will be critical to incorporate data from non-bilaterian animals such as sponges and ctenophores (Dunn et al. 2015). Non-bilaterians are more phylogenetically divergent from each other, and from other animals, than are the research models listed above (Dohrmann and Wörheide 2017; Schuster et al. 2017), but detailed mechanistic knowledge about their cell and developmental biology is limited. However, progress in this area has accelerated due to advances in comparative genomics, with the initially surprising result that starkly different organisms share highly conserved developmental regulatory genes (McGinnis et al. 1984; Kusserow et al. 2005; Technau et al. 2005; Nichols et al. 2006). An important part of understanding animal body plan evolution will be to explain how conserved bilaterian regulatory genes/pathways function in non-bilaterian animals, how they may have functioned ancestrally, and how changes to their functions may have contributed to morphological evolution.

To begin to address these questions, the pleiotropic gene β-catenin is of considerable interest because it functions in developmentally important processes, including cell adhesion and transcriptional regulation in the Wnt/β-catenin signaling pathway. In cell adhesion, β-catenin interacts with the cytoplasmic tail of cadherin receptors and with α-catenin, an unrelated protein that links the adhesion complex to the actin cytoskeleton (Pokutta and Weis 2007; Shapiro and Weis 2009). This molecular complex forms the foundation of the adherens junction (AJ) and serves as the primary cell-cell adhesion mechanism in bilaterian tissues, including epithelia where the AJ also serves as a spatial cue for the establishment of tissue polarity (Capaldo and Macara 2007). In Wnt/β-catenin signaling, β-catenin is a downstream effector that translocates to the nucleus in response to Wnt signals, where it forms a complex with the transcription factor TCF/Lef. Wnt/β-catenin signaling regulates hundreds of downstream target genes and is involved in developmental processes ranging from stem cell maintenance and renewal to axial patterning and gastrulation (Clevers and Nusse 2012).

An early study of β-catenin evolution in the sea anemone *Nematostella vectensis* predicted that the dual signaling and adhesion functions of β-catenin were ancient, and already established in early animal ancestors (Schneider et al. 2003). Consistent with this view, it has since been established that not only β-catenin, but most components of the AJ and the Wnt/β-catenin pathway are conserved in other non-bilaterian lineages as well. Moreover, expression studies indicate roles for the Wnt/β-catenin pathway in developmental patterning [reviewed in (Holstein 2012)] in cnidarians (Hobmayer et al. 2000; Wikramanayake et al. 2003; Lee et al. 2006; Momose et al. 2008), ctenophores (Pang et al. 2010; Jager et al. 2013) and sponges (Srivastava et al. 2010; Leininger et al. 2014), and biochemical data indicate that the cadherin/catenin adhesion complex has conserved roles in epithelial cell adhesion in *N. vectensis* (Clarke et al. 2016).

Among non-bilaterian animals, studies of β-catenin in sponges are of particular interest because of their divergent body plan and developmental differences. Whereas axial patterning and gastrulation are hypothesized to be ancestral functions for Wnt/β-catenin signaling (Kusserow et al. 2005; Lee et al. 2006), the adult sponge body plan lacks clear axial polarity and it is contentious as to whether they undergo gastrulation (Nakanishi et al. 2014). With respect to a possible role for β-catenin in cadherin-based adhesion in sponges, ultrastructural studies indicate that their tissues generally lack prominent AJs (Leys et al. 2009). Instead, sponge cell adhesion (at least in demosponges) has long been attributed to a secreted glycoprotein complex called the Aggregation Factor (Henkart et al. 1973; Müller and Zahn 1973; Fernàndez-Busquets and Burger 1997; Grice et al. 2017), contributing to the view that sponge tissue organization is fundamentally different from the organization of epithelia in other animals.

Several studies have begun to characterize β-catenin function in sponges by examining its expression dynamics. In the demosponge *Amphimedon queenslandica* and in the calcareous sponge *Sycon ciliatum*, β-catenin and other Wnt pathway components are differentially expressed during embryonic development, indicating a conserved role in patterning (Srivastava et al. 2010; Leininger et al. 2014). In adult tissues of *S. ciliatum*, β-catenin is also expressed in the choanoderm (feeding epithelium) and in a ring of migratory cells around the osculum (exhalant canal) (Leininger et al. 2014). Other studies have taken a pharmacological approach to study the function of Wnt/β-catenin signaling by treating the sponges with Glycogen Synthase Kinase 3 Beta (GSK3B) inhibitors (Lapébie et al. 2009; Windsor and Leys 2010). GSK3B is a negative regulator of Wnt signaling that functions by phosphorylating free cytosolic β-catenin, leading to its ubiquitination and degradation by the proteasome. When GSK3B is inhibited (i.e., in the presence of Wnt ligands), cytosolic β-catenin accumulates, translocates to the nucleus and activates TCF/Lef and the transcription of Wnt target genes. In the sponge *Oscarella lobularis*, GSK3B inhibitors caused the formation of ectopic ostia (incurrent water pores) and caused morphological changes to the exopinacoderm (outer epithelium) (Lapébie et al. 2009). In the freshwater sponge *Ephydatia muelleri*, GSK3B inhibitors caused the formation of ectopic oscula (exhalant water canals) and malformed choanocyte chambers (filter feeding structures) (Windsor and Leys 2010).

With respect to a possible role for β-catenin in cadherin-based adhesion in sponges, all AJ components have been identified (Srivastava et al. 2010; Nichols et al. 2012; Riesgo et al. 2014), but experimental evidence for their function is limited. Sequence analyses and structural predictions indicate that sponge homologs of β-catenin and classical cadherins are sufficiently similar to their bilaterian counterparts to undergo conserved protein-protein interactions. From an experimental perspective, a single study of the homoloscleromorph sponge *Oscarella pearsei* [previously called O. *carmela* (Ereskovsky et al. 2017)] detected the interaction between β-catenin and a classical cadherin by yeast two-hybrid screen (Nichols et al. 2012).

In general, hypotheses about early animal evolution rest heavily on assumptions about developmental regulatory genes in organisms where their function and interactions have been insufficiently tested. In the case of β-catenin, reported gene expression patterns may reflect the conservation of one or more bilaterian functions, or entirely alternative functions; it is impossible to distinguish between these possibilities from expression data alone. Likewise, phenotypes resulting from pharmacological inhibition of GSK3B may not reflect perturbations to the Wnt/β-catenin pathway; GSK3B is pleiotropic, and β-catenin is not a confirmed GSK3B substrate in sponges. Here, we examine β-catenin function in the freshwater sponge, *E. muelleri*. Using a polyclonal antibody raised against *E. muelleri* β-catenin (*Em* β-catenin), we 1) identify endogenous *Em* β-catenin binding partners; 2) characterize discrete subcellular localization patterns of *Em* β-catenin at cell contacts (cell-cell adhesion), the nucleus (signaling), and at possible cell-substrate adhesions in the attachment epithelium (novel function?), and; 3) find that *Em* β-catenin is a probable substrate of GSK3B.

## RESULTS

### Adherens Junction (AJ) and Wnt pathway components in the *E. muelleri* transcriptome

We searched the published *E. muelleri* transcriptome (Peña et al. 2016) and detected a single ortholog of β-catenin. The *Em β-catenin* transcript has a predicted 2,685 bp coding sequence, corresponding to an 895 aa predicted protein. Furthermore, as reported from multiple other sponge species (Nichols et al. 2006; Adamska et al. 2010; Fahey and Degnan 2010; Leininger et al. 2014; Riesgo et al. 2014), we detected conserved homologs of AJ and Wnt signaling pathway components (Supplement Table S1).

*Em* β-catenin has 53% sequence identity to *Mus musculus* β-catenin, 50% identity to *Drosophila melanogaster*, and between 30-93% identity with β-catenin from other sponge species. The core of the *Em* β-catenin protein is predicted to contain 12 armadillo (Arm) repeats flanked by less conserved, and presumably unstructured N- and C-terminal regions. In bilaterians, the Arm repeat region is known to serve as the binding interface for interacting with APC, Axin and TCF/Lef which are involved in the Wnt signaling pathway, but also classical cadherins, a component of the AJ (Valenta et al. 2012). By aligning *Em* β-catenin to structurally and functionally characterized orthologs in bilaterians, we found the conservation of residues Lys312 and Lys435 which are required for binding of mouse β-catenin to E-cadherin (Huber and Weis 2001) (see Fig1 and Supplement Fig S1). Also, *Em* β-catenin has conserved GSK3B and CKI phosphorylation sites in the N-terminal domain (Ser33, 37, Thr41, Ser45 in mouse), which are required for initiating the “destruction complex” in the Wnt signaling pathway (see Fig1 and Supplement Fig S1). The binding motif for α-catenin (a constitutive AJ component) in the N-terminal domain was found to be conserved to a lesser extent (52% sequence identity with mouse) and only 2 of 5 structurally important residues, but 7 out of 10 binding surface residues (Miller et al. 2013) were conserved in *Em* β-catenin (Supplement Fig S1).

**Figure 1.**
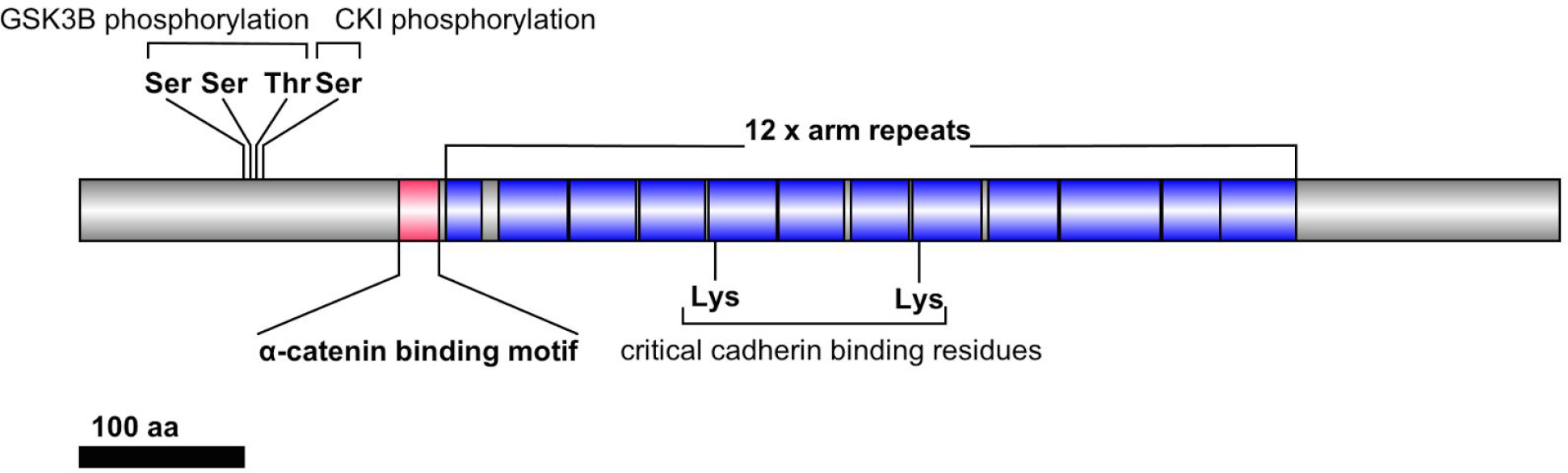
Predicted domain architecture of *Em* β-catenin. The protein core contains 12 predicted Armadillo (Arm) repeats flanked by N- and C-terminal regions. The N-terminus has conserved GSK3B and CKI phosphorylation residues and a putative α-catenin binding site. Critical cadherin binding residues are also conserved. The sequence and predicted structural conservation of this protein suggests plausibly conserved roles in Wnt signaling and at the Adherens Junction (AJ).

### Adherens Junction proteins co-precipitate with *Em* β-catenin

The discrete signaling and adhesion functions of β-catenin are defined by interactions with different proteins. To identify endogenous *Em* β-catenin binding-partners, we raised an antibody against a 24 kDa recombinant protein corresponding to the N-terminal region. The resulting antibody was affinity purified and its specificity was tested by Western Blot against denatured whole-cell lysates. As apparent in Fig 2A, a predominant band of the expected size (~110 kDa) was detected, but lower molecular weight bands were also sometimes present. These lower molecular weight bands could be minimized by reducing handling time during preparation of lysates, and sometimes only a single band was detected (see Supplement Fig S3), supporting their interpretation as *Em* β-catenin degradation products. To further validate this claim, we preadsorbed the antibody with recombinant antigen prior to blotting and all bands were eliminated (Supplement Fig S4).

**Figure 2.**
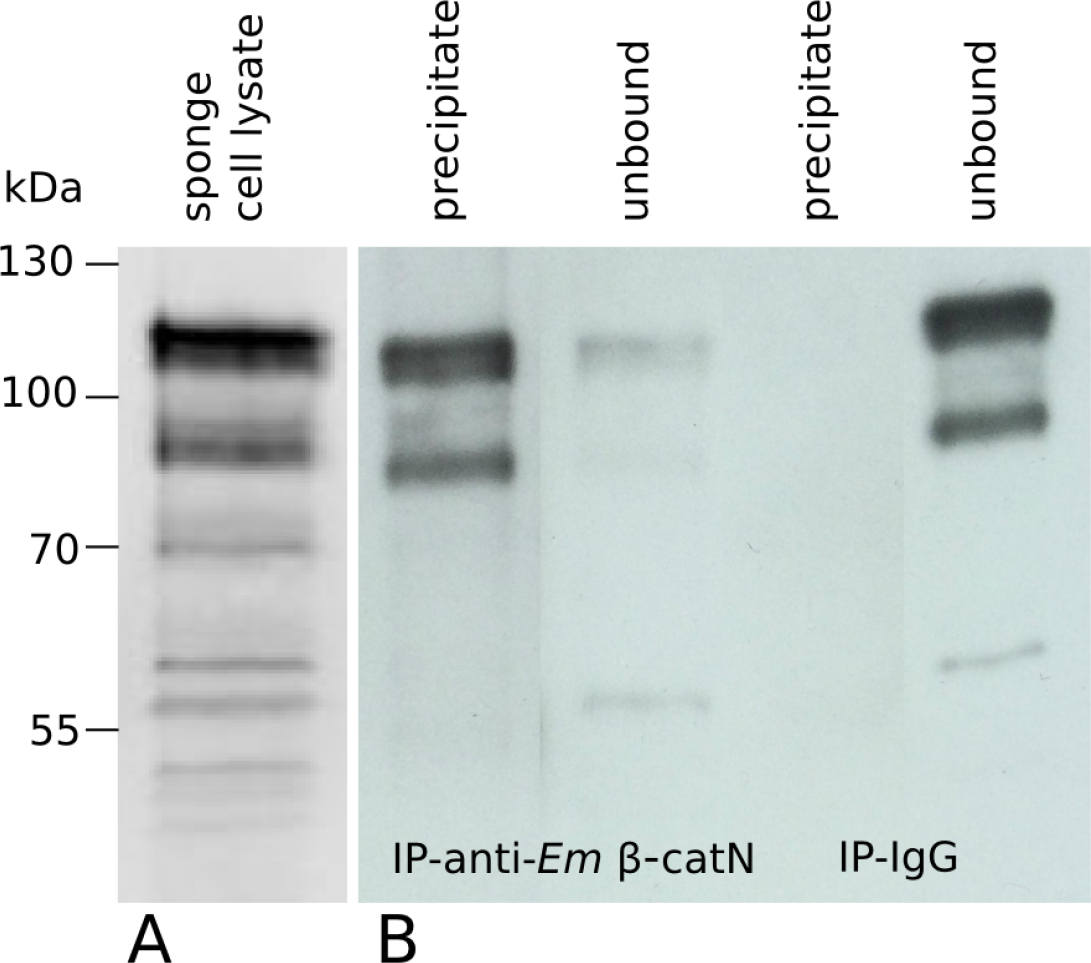
*Em* β-catenin antibody specificity. The specificity of the *Em* β-catenin antibody (anti-Em β-catN) was tested by Western Blot against denatured whole-cell lysates of *E. muelleri* (A). A predominant band of the expected size (~110 kDa) was detected, but lower molecular weight bands were also sometimes present and interpreted as degradation products (see Fig S3 and S4 for further details). The antibody was further validated by immunoprecipitation (IP) and was compared to IP with an IgG control. The precipitate and unbound (i.e. flowthrough) fractions were analyzed by Western Blot (B). The lanes with precipitates on the blot show that *Em* β-catenin is being pulled down by the anti-Em β-catN antibody, but not by the IgG control.

To validate the specificity of antibody under non-denaturing conditions, we performed immunoprecipitation (Fig 2B) coupled with mass spectrometry. As summarized in Table 1 (full results available in Supplement), the most abundantly detected peptides in the precipitate corresponded to *Em* β-catenin. Two co-precipitates were α-catenin (*Em* α-catenin) and a classical cadherin [*Em* CDH2, see (Peña et al. 2016)]; both are homologs of AJ components. Another co-precipitate was coiled-coil domain-containing protein 91 (ccdc91, also known as p56). The only available studies of this protein in animals reveal that it is a trans-Golgi accessory protein involved in vesicle transport between the Golgi and the lysosome (Mardones et al. 2007). There is no precedent in the literature for interactions between ccdc91 and β-catenin. We found no evidence for conserved interactions with Wnt pathway components, such as TCF/Lef, Axin or APC.

**Table 1.** Mass spectrometry results of Co-Immunoprecipitation

| Identity | T-PEP ID | Summed # unique peptides | Summed # total spectrum | MW (kDa) |
| --- | --- | --- | --- | --- |
| ***Em* β-catenin** | m.48575 | 32 | 104 | 98 |
| **ccdc-91/p56** | m.42889 | 14 | 22 | 79 |
| ***Em* CDH2** | m.5230 /<br>m.46898 | 10 | 12 | 587 |
| ***Em* α-catenin** | m.75406 | 5 | 11 | 102 |

### Distinct *Em* β-catenin populations at cell contacts (adhesion) and in the nucleus (signaling)

Functionally distinct β-catenin populations exhibit different subcellular localization patterns; adhesion-related populations localize to cell-cell contacts, whereas signaling populations localize to the nucleus. To further test the signaling and adhesion roles of *Em* β-catenin we performed immunostaining on somatic tissues in gemmule hatched juveniles. One tissue, the apical endopinacoderm (AEP), is an epithelium cored by tracts of F-actin that form dense plaques at points of alignment between neighboring cells (De Ceccatty 1986; Elliott and Leys 2007) that resemble an AJ. By immunostaining with anti-Em β-catenin we detected staining at F-actin plaques in the AEP (Fig 3). This result is consistent with detected interactions between *Em* β-catenin with *Em* α-catenin and *Em* CDH2, and indicates a conserved role for cadherin-mediated adhesion within the AEP.

**Figure 3.**
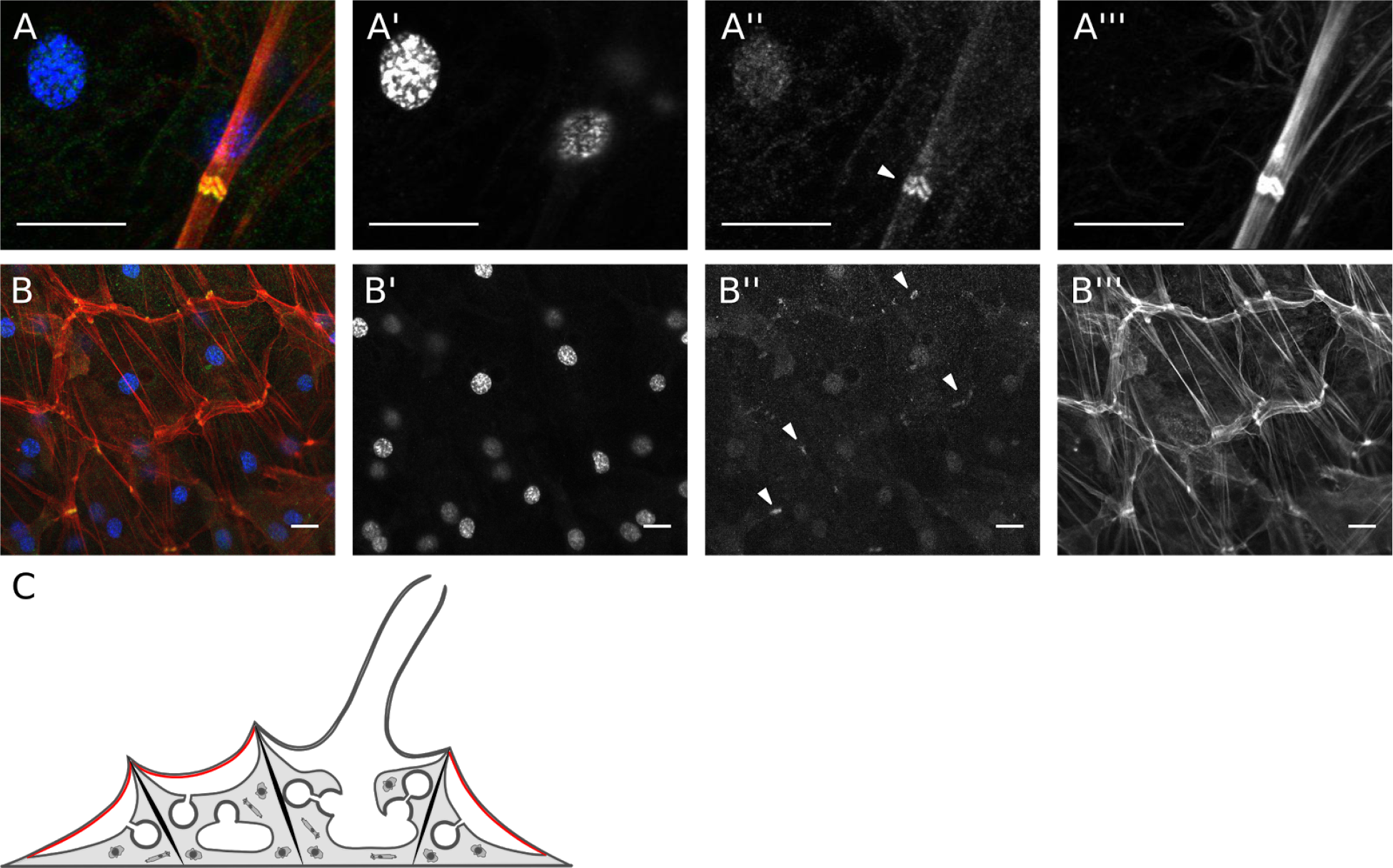
*Em* β-catenin co-localizes with plaques of F-actin at cell contacts in the apical endopinacoderm (AEP). The AEP is an epithelium cored by tracts of F-actin that form dense plaques at points of alignment between neighboring cells. (A) Em β-catenin localizes to these plaques (arrowheads), consistent with evidence that the AEP is the contractile epithelium. (B) Widefield view showing general organization of F-actin tracts and plaques in the AEP. Composite images (A/B) with DNA (A’/B’) in blue, anti-Em β-catN (A”/B”) in green, and F-actin (A’”/B’”) in red. Scale bar: 10 µm. (C) Illustrative cross section of *E. muelleri* with AEP in red.

In the attachment epithelium, a cellular monolayer at the interface with the substrate, we detected continuous cell-boundary staining of *Em* β-catenin, consistent with the presence of a belt AJ. This staining was relatively low intensity (see Fig 4), but consistently higher than background staining levels in secondary-only negative controls, and is eliminated by preadsorption of the antibody with recombinant antigen (Supplement Fig S5). More pronounced *Em* β-catenin staining was evident in the cytosol, at the ends of F-actin stress fibers, which differed from actin tracts of the AEP in that they were not regularly oriented in the cell, they were not aligned between neighboring cells, and did not extend to the full diameter of any individual cell (Fig 4A; Supplement Fig S6). Superficially, these structures resemble focal adhesions (FAs), but *Em* β-catenin is localized where FA proteins such as talin, paxillin, and integrins would be expected in FAs. This result is surprising because, like FAs in bilaterians, these structures have previously been interpreted as cell-substrate adhesions (De Ceccatty 1986; Leys and Hill 2012), whereas β-catenin is not typically involved in cell-substrate adhesion in other animals.

**Figure 4.**
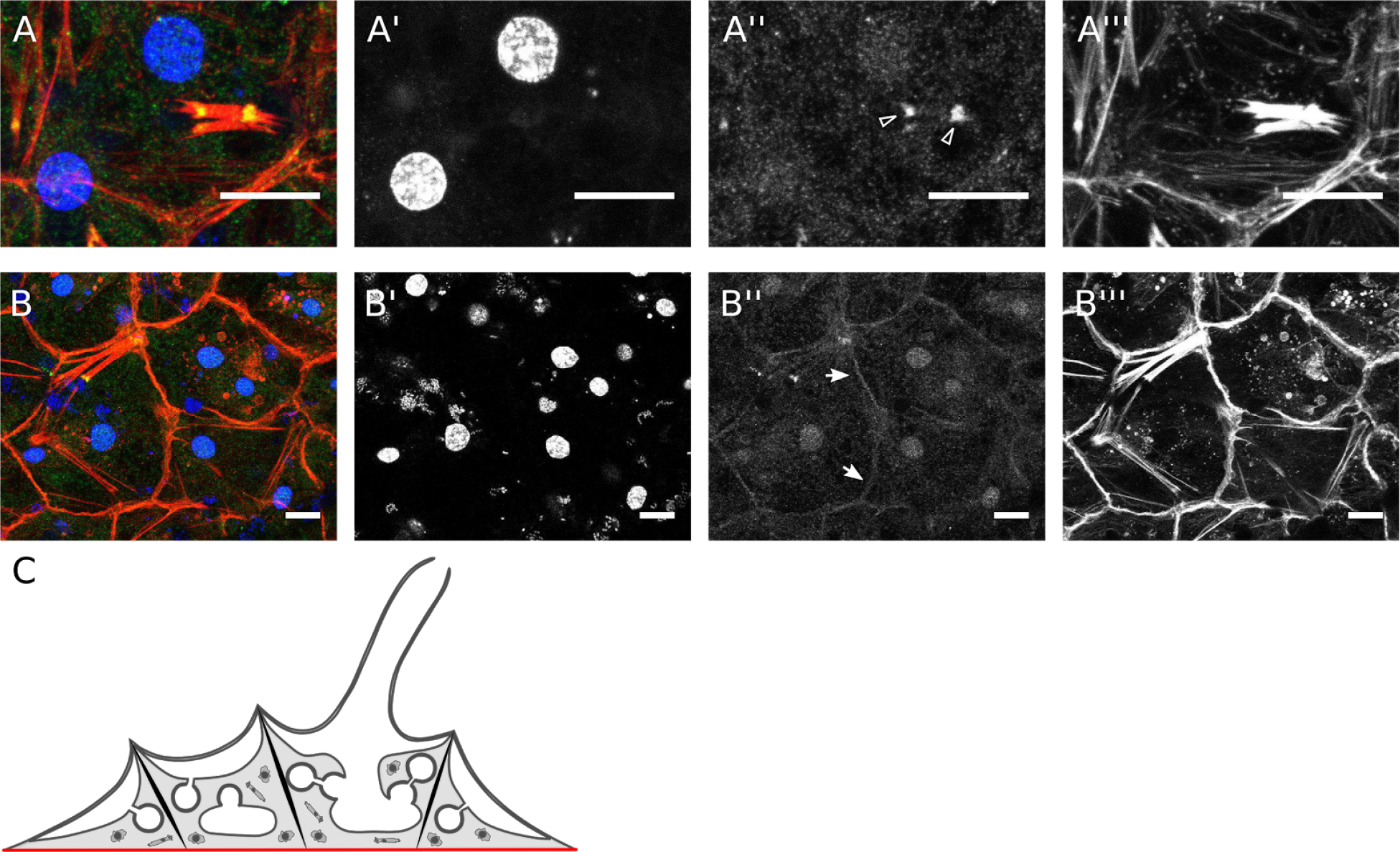
*Em* β-catenin localization in the attachment epithelium at cell boundaries and at apparent cell-substrate adhesion structures. The attachment epithelium is a cellular monolayer at the interface with the substrate. (A) Em β-catenin was enriched (arrowheads) at the distal ends of F-actin stress fibers that differ from F-actin tracts in the AEP (described in the text) and superficially resemble focal adhesion. Fig S6 shows an example of the frequency and orientation of these structures in this tissue. (B) *Em* β-catenin was also detected as a continuous cell-boundary staining (arrows) in this tissue, which is consistent with the presence of a belt AJ. Composite images (A) with DNA (A’) in blue, anti-Em β-catN (A”) in green, and F-actin (A’”) in red. Scale bar: 10 µm. (B) Illustrative cross section of *E. muelleri* with attachment epithelium in red.

We might also expect to find cell-substrate adhesions in highly migratory cells of the mesohyl, such as archeocytes, as the dynamic regulation of FAs is important for cell migration in some well-studied bilaterian models (Mayor and Etienne-Manneville 2016). However, we find no evidence for cell-substrate adhesions in archeocytes, nor are there filopodia-associated *Em* β-catenin populations in these cells (Fig 5).

**Figure 5.**
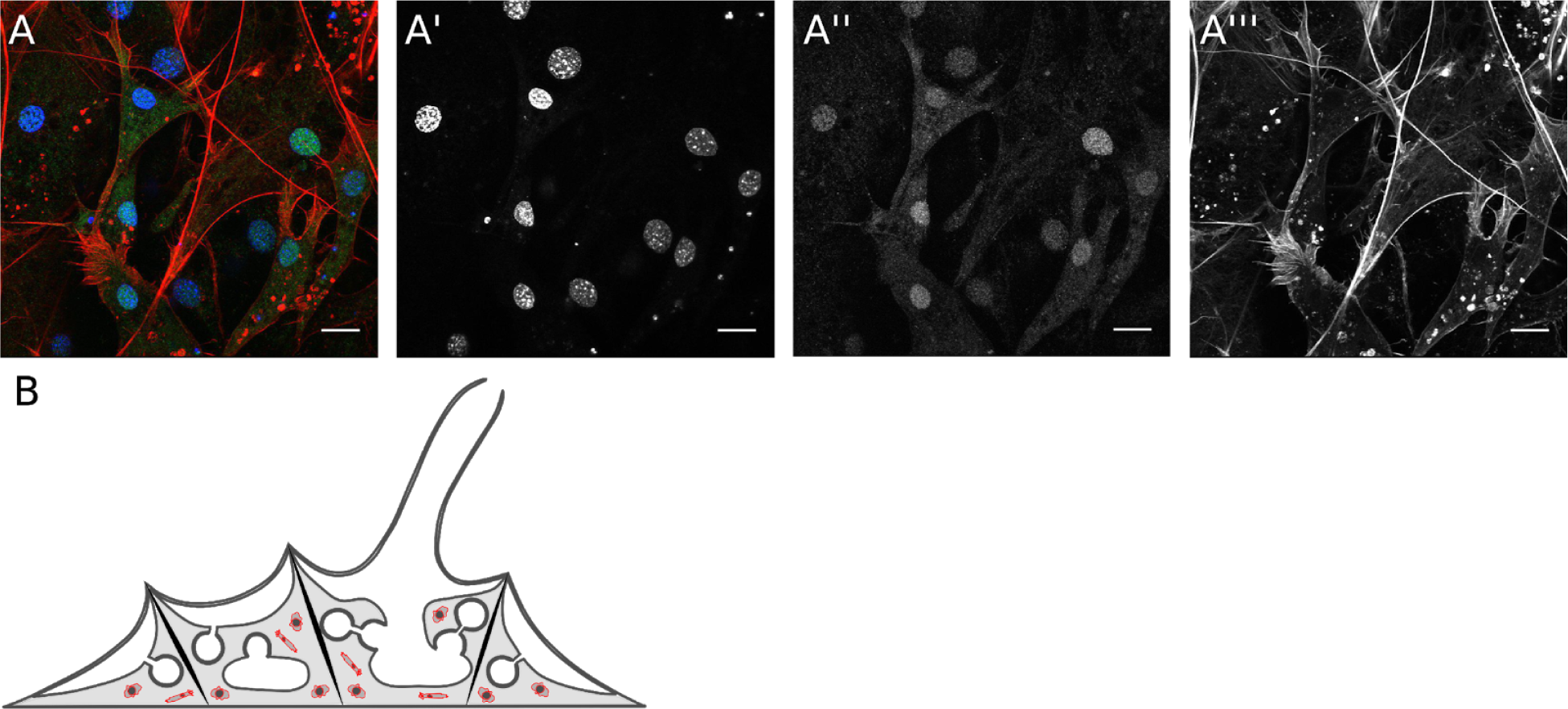
Nuclear localization of *Em* β-catenin in migratory cells in the mesohyl. The mesohyl is a collagenous extracellular matrix populated by migratory cells. These migratory cells include archeocytes, which are considered to have stem-cell like properties. The nuclei in these migratory cells in the mesohyl (panel A) show enrichment for *Em* β-catenin. Composite images (A) with DNA (A’) in blue, anti-Em β-catN (A”) in green, and F-actin (A’”) in red. Scale bar: 10 µm. (B) Illustrative cross section of *E. muelleri* with migratory cells in mesohyl in red.

Another distinctive, polarized epithelium in sponges is the choanoderm, which consists of spherical chambers lined by collar cells (i.e. choanocytes), each with an apical ring of microvilli that surrounds a single flagellum. These cells pump water through internal canals and phagocytize bacterial prey. As in the attachment epithelium, newly developing choanocyte chambers (see Fig 6A) exhibited continuous cell boundary staining of *Em* β-catenin. However, in mature choanocyte chambers (see Fig 6B and C), this staining pattern was less evident, and instead *Em* β-catenin was enriched at points of contact between three adjacent cells (Fig 6C).

**Figure 6.**
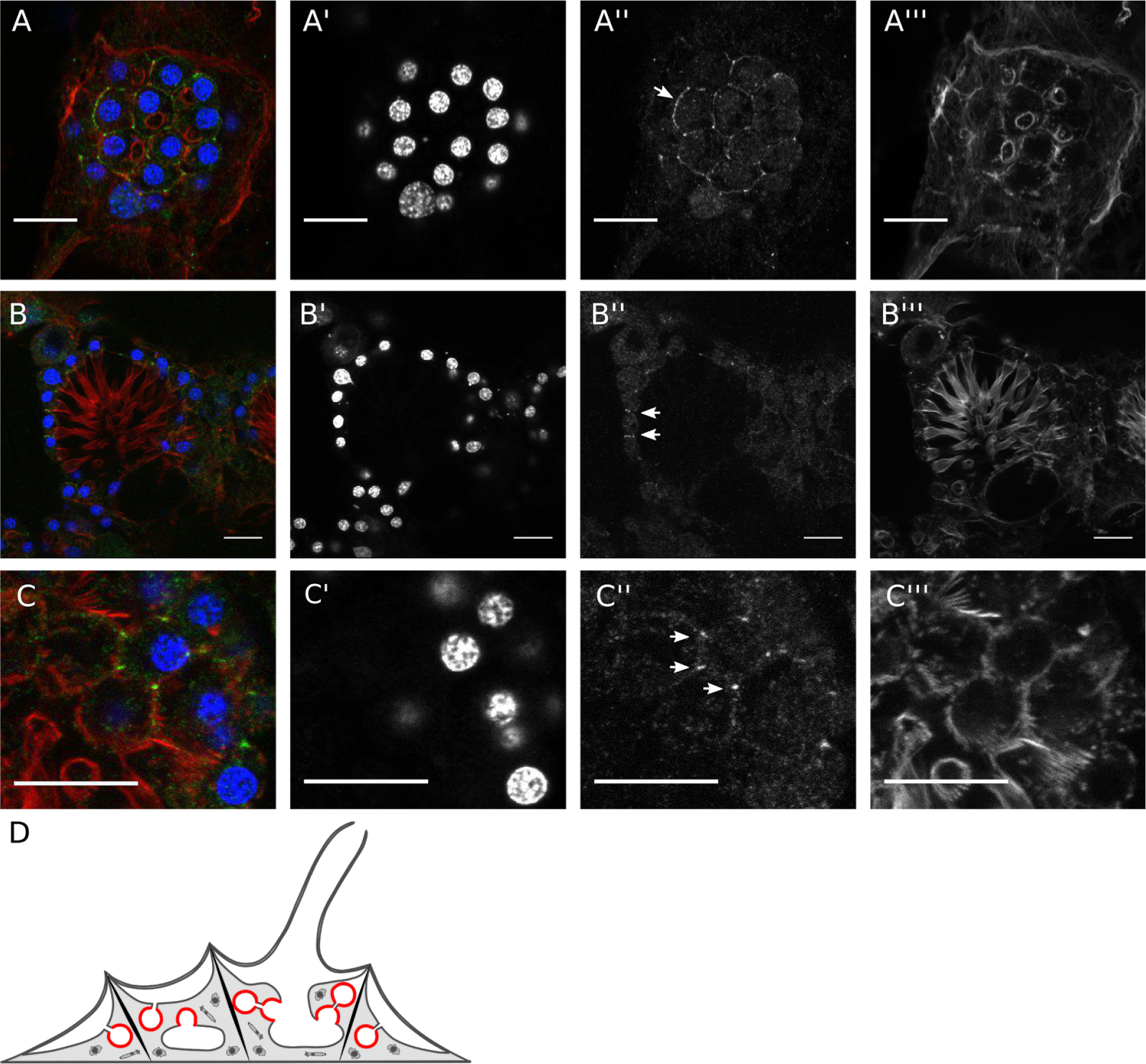
*Em* β-catenin localizes to cell boundaries in the choanoderm. The choanoderm is the internal feeding epithelium and consists of spherical chambers lined by collar cells (i.e. choanocytes). In newly developing choanocyte chambers (panel A), Em β-catenin is detected as a continuous cell boundary staining (arrow), just as in the attachment epithelium (Fig 4B). However, in mature choanocyte chambers (panel B and C) continuous cell boundary staining was less clear and Em β-catenin was enriched at points of contact between three adjacent cells (arrow). Composite images (A/B/C) with DNA (A’/B’/C’) in blue, anti-Em β-catN (A”/B”/C”) in green, and F-actin (A’”/B’”/C’”) in red. Scale bar: 10 µm. (D) Illustrative cross section of *E. muelleri* with choanoderm in red.

Whereas we did not find evidence of conserved interactions between *Em* β-catenin and other Wnt pathway components by immunoprecipitation, we further examined the possible signaling roles of *Em* β-catenin by immunostaining to test for nuclear populations. We detected nuclear *Em* β-catenin in the attachment epithelium (Fig 7) as well as in migratory cells in the mesohyl (Fig 5), which is consistent with a conserved role for *Em* β-catenin as a transcriptional coactivator of TCF/Lef. We did not detect nuclear *Em* β-catenin in choanocytes (Fig 6). There may be nuclear staining in the AEP as well, but the dermal surface of the sponge is composed of two thin epithelia that surround a thin extracellular matrix populated by migratory cells (Elliott and Leys 2007; Adams et al. 2010). Because these tissues are thin, closely spaced and irregularly shaped relative to the plane of focus, it is difficult to resolve whether *Em* β-catenin-positive nuclei are associated with epithelial tissue or migratory cells in this region.

**Figure 7.**
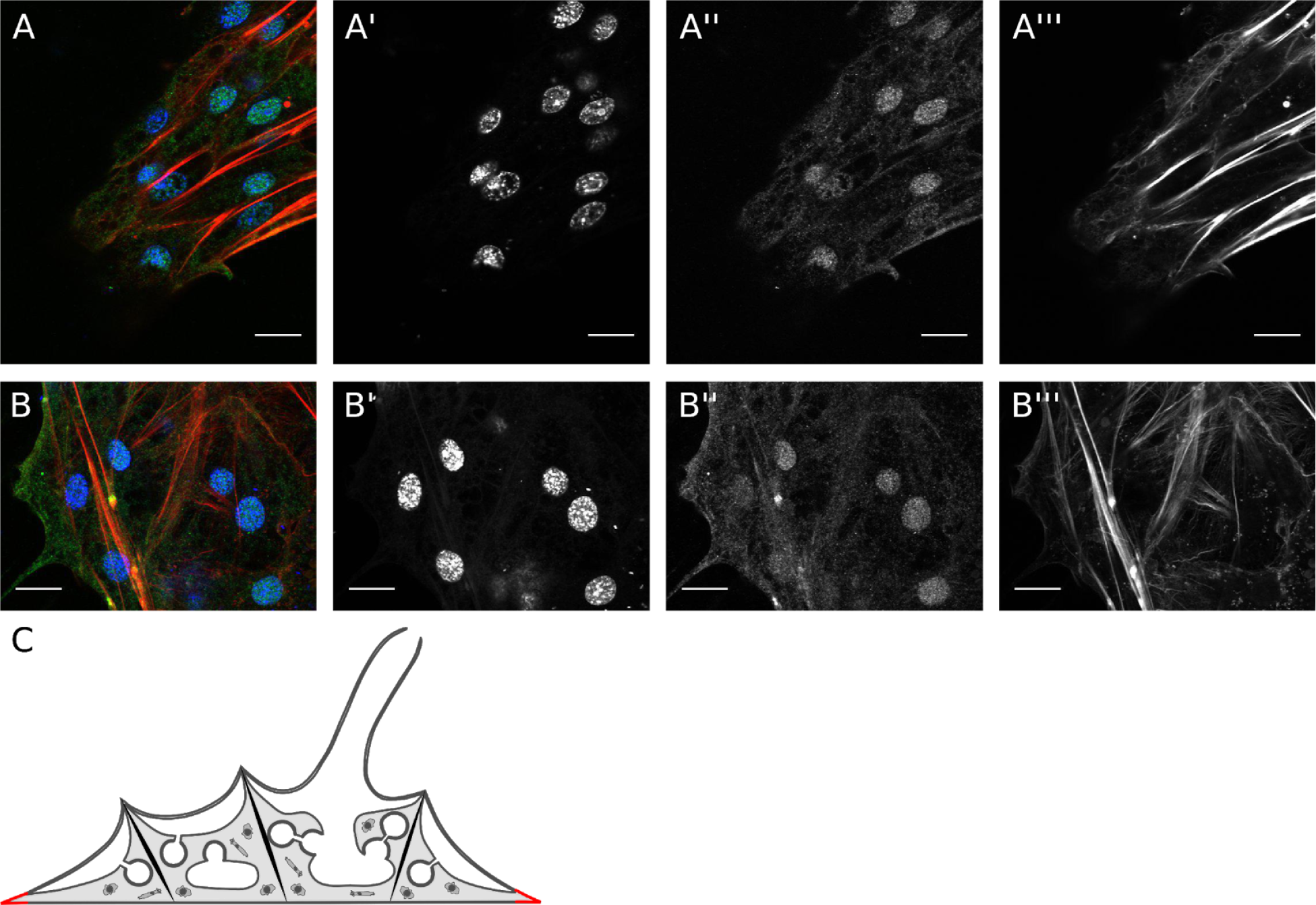
Nuclear localization of *Em* β-catenin in cells of the leading edge of the attachment epithelium. Sponges are capable of entire-colony mobility and in juveniles of *E. muelleri* this results in collective migration of the attachment epithelium. Nuclear localization of *Em* β-catenin can be found throughout the whole attachment epithelium (Fig 4B), but is most evident in cells of the leading edge of the migrating attachment epithelium (panel A and B). Composite images (A/B) with DNA (A’/B’) in blue, anti-Em β-catN (A”/B”) in green, and F-actin (A’”/B’”) in red. Scale bar: 10 µm. (C) Illustrative cross section of *E. muelleri* with leading edge of the attachment epithelium in red.

### *Em* β-catenin is a conserved substrate of GSK3B

GSK3B is a negative regulator of β-catenin in the Wnt/β-catenin signaling pathway, and previous studies have shown that treating sponges with multiple, independent GSK3B inhibitors has reproducible phenotypic effects (Lapébie et al. 2009; Windsor and Leys 2010). To more directly test if the effects of GSK3B are related to Wnt/β-catenin signaling we examined whether GSK3B inhibitors affect endogenous *Em* β-catenin phosphorylation states. We treated juvenile sponges with the GSK3B inhibitors Alsterpaullone (AP) and BIO, and monitored their effects by Western Blot (Fig 8). Typically, when GSK3B is active, β-catenin is being phosphorylated and targeted for degradation. When GSK3B is inhibited, β-catenin is no longer phosphorylated and cytosolic β-catenin accumulates and translocates to the nucleus where it activates the Wnt pathway. We detected an increase in electrophoretic mobility of *Em* β-catenin in lysates from GSK3B inhibited sponges, which we interpret as a change in the *Em* β-catenin phosphorylation state (dephosphorylated *Em* β-catenin is expected to migrate faster in the gel). As confirmation, the electrophoretic migration rate of *Em* β-catenin was also monitored in sponge lysates that were variously treated with phosphatases, phosphatase inhibitors, both, or neither. We found that phosphatase treatment increased the electrophoretic mobility of *Em* β-catenin, whereas phosphatase inhibition decreased its electrophoretic mobility (Supplement Fig S7).

**Figure 8.**
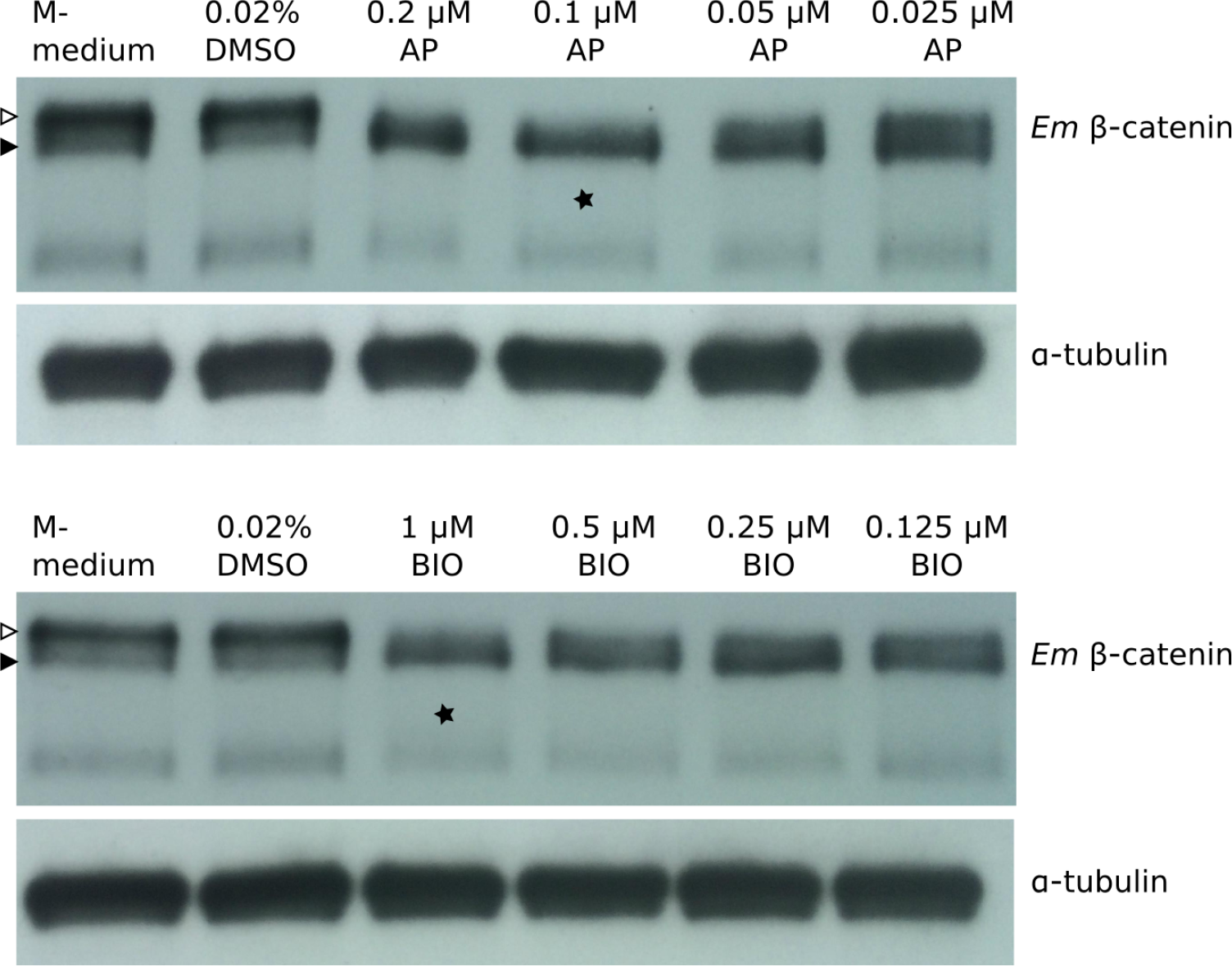
Western blot showing effect of AP and BIO on phosphorylated *Em* β-catenin protein levels from sponge cell lysate. Exposing sponges to GSK3B inhibitors results in a shift in electrophoretic mobility of *Em* β-catenin. When GSK3B is inhibited, *Em* β-catenin is no longer phosphorylated and migrates faster on the gel [closed arrowhead: active *Em* β-catenin (i.e. dephosphorylated)]. When GSK3B is active, *Em* β-catenin is phosphorylated and migrates slower on the gel (open arrowhead: GSK3B - phosphorylated *Em* β-catenin). To find the optimal effective dosage of GSK3B inhibitors, we tested a range of concentrations and found that 0.1 µM AP and 1 µM BIO (asterisk) results in a complete shift from phosphorylated to active *Em* β-catenin. The loading control a-tubulin confirms that concentrations of cell lysate were loaded in each lane.

Through titration of the GSK3B inhibitors we determined that treatment with ~ 0.1 µM AP or ~1 µM BIO resulted in a complete shift from phosphorylated to dephosphorylated states of *Em* β-catenin and was therefore considered the optimal effective dosage (Fig 8). Gemmules treated with this dosage developed into sponges with dense populations of cells that could not easily be correlated to differentiated tissues, but canals were clearly lacking and the choanoderm was absent (Fig 9). Thus, GSK3B inhibition at the optimal effective dosage disrupts the development of the aquiferous system, including differentiation of the choanoderm. These changes in morphology as a result of treating with GSK3B inhibitors correspond to previously reported phenotypes that were characterized in greater detail by light and electron microscopy (Windsor and Leys 2010).

**Figure 9.**
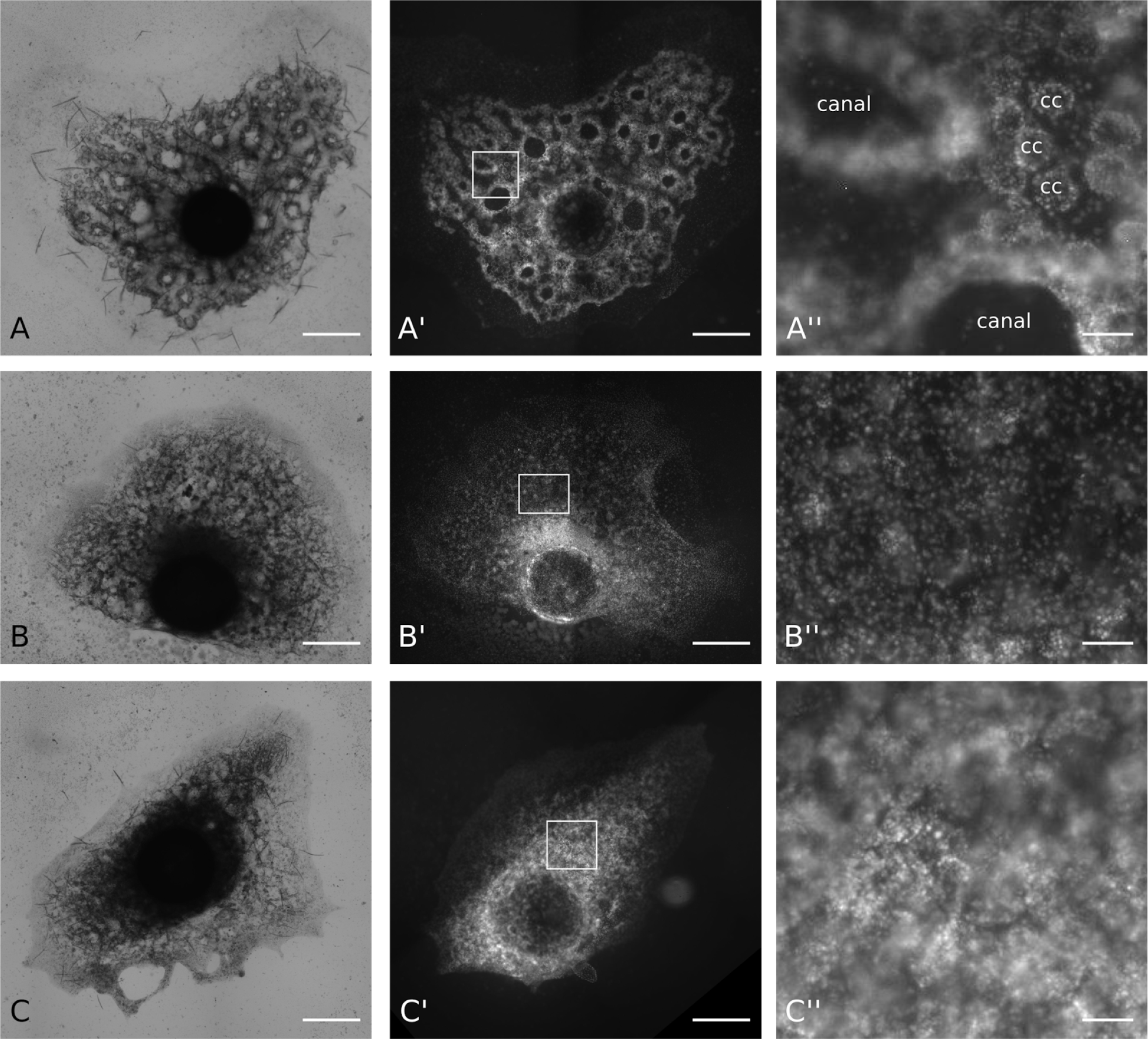
Phenotypic effects of GSK3B inhibitors. Sponges that were exposed to the optimal effective dosage 0.1 µM AP and 1 µM BIO were assumed to possess only active β-catenin, and therefore have an activated Wnt/β-catenin signaling pathway. Sponges treated with dosage (B and C) exhibited dense populations of cells and they clearly lacked canals and the choanoderm was either absent or malformed. For comparison, the control treatment (A) clearly shows canals and choanocyte chambers. (A/B/C) Brightfield images, (A’/B’/C’) Fluorescence images with DAPI staining showing DNA, (A”/B”/C”) magnification of Fluorescence image to show the canals (canal) and choanocyte chambers (cc) in more detail. Scale bar left and middle panel: 500 µm, right panel: 50 µm.

Efforts were also made to directly disrupt *Em* β-catenin function using RNAi and Vivo-morpholinos. These techniques caused non-specific phenotypic effects (which varied between individuals collected from different localities), but no measurable change in *Em* β-catenin levels as measured by Western Blot (Supplement Fig S8). There are no well-established techniques for functional studies in sponges, so reporting the experimental detail of these negative trials is relevant to aid in future technique development.

## DISCUSSION

β-catenin is thought to have played a critical role in animal body plan evolution due to its pleiotropic roles in development signaling (Wnt signaling pathway) and in cell-cell adhesion (the Adherens Junction). These roles are broadly conserved in all studied animals, but functional data are available from only cnidarians and bilaterians. Earlier diverging animal lineages, including sponges, have conserved homologs of Wnt pathway and cell adhesion genes but there is limited experimental evidence pertaining to their function. Thus, hypotheses about the role of Wnt signaling and cadherin-based cell adhesion during the earliest periods of animal evolution depend partly upon improved understanding of β-catenin function in these understudied lineages.

Looking outside of animals, there is evidence for a possible β-catenin homolog (*Aardvark*) in the social amoeba *Dictyostelium discoideum* (Grimson et al. 2000) that interacts with an α-catenin-like protein to regulate the polarity of the tip epithelium of the fruiting body (Dickinson, Nelson, et al. 2011). Nevertheless, D. *discoideum* lacks both cadherins and Wnt pathway components. If we consider lineages that are more closely related to animals, cadherins have been detected in several unicellular opisthokonts, but they lack β-catenin interacting regions, they lack conserved β-catenin orthologs, and there is no evidence for other conserved Wnt pathway components (Abedin and King 2008; Nichols et al. 2012). Collectively, these data suggest that the pleiotropic functions of β-catenin in cell adhesion and Wnt signaling are an early innovation in the animal stem lineage.

### The Adherens Junction in sponge tissues

Sponges have been predicted to have an AJ based upon genomic and transcriptomic data from diverse species (Nichols et al. 2006; Srivastava et al. 2010; Nichols et al. 2012; Riesgo et al. 2014), and upon limited ultrastructural evidence [reviewed in (Leys et al. 2009)]. Our *Em* β-catenin immunoprecipitation data provide new experimental support for endogenous interactions between AJ components in *E. muelleri*. Notably, we detected an interaction between *Em* β-catenin with *Em* CDH2 and *Em* α-catenin, even though the residues required for the latter interaction in vertebrates (Miller et al. 2013) are incompletely conserved in *E. muelleri*. This result underscores the limitations of functional predictions made using bioinformatic approaches [elaborated in (Dickinson, Weis, et al. 2011)], and parallels the example of Axin in the sponge *O. pearsei*, which lacks the expected binding motif for β-catenin but was identified as a binding partner via yeast two-hybrid screen (Nichols et al. 2012). A possible explanation for why *Em* CDH2 and not *Em* CDH1 was detected as a candidate *Em* β-catenin binding partner is that *Em* CDH1 may be expressed at a different developmental stage than the juvenile tissues examined. In bilaterians, classical cadherins often exhibit tissue-specific expression. For example, in mammals, E-cadherin is expressed in epithelial tissues, N-cadherin expressed in the nervous system, and P-cadherin in the placenta (Yoshida-Noro et al. 1984; Hatta and Takeichi 1986; Nose and Takeichi 1986; Takeichi 1988). As sponges appear to be largely epithelial organisms it is intriguing to consider that they may have different adhesion receptors expressed in different contexts. Spatiotemporal differences in expression of cadherin paralogs may reveal differences in the evolutionary history and organization of different sponge tissues.

Immunostaining of *Em* β-catenin further supports the existence of the AJ in sponge tissues. Continuous cell boundary staining is evident in the attachment epithelium (Fig 4) as well as in newly developing choanocyte chambers (Fig 6A). This staining pattern resembles the organization of the AJ in cnidarian and bilaterian epithelial tissues, which form a continuous ‘belt’ around the periphery of the cell, thereby dividing the plasma membrane into apical and basolateral domains. Together with occluding junctions (tight junctions in vertebrates and septate junctions in invertebrates), the AJ can help create a barrier that regulates paracellular transport, effectively separating the environment outside of the tissue from the interior environment – a fundamental feature of animal multicellular organization (Banerjee et al. 2006; Hartsock and Nelson 2008; Zihni et al. 2016). There is evidence that freshwater sponges have “sealed” epithelia (Adams et al. 2010), but it is not yet established which molecules are involved.

Instead of continuous cell-boundary staining, mature choanoderm tissue (Fig 6C) and the AEP (Fig 3) were found to have probable adhesion-associated *Em* β catenin populations with a much more restricted distribution. In mature choanocyte chambers, *Em* β-catenin was largely enriched at points of contact where three neighboring cells meet; this localization pattern superficially resembles tricellular junctions of *Drosophila*, although they have a different molecular composition than the AJ. In the AEP, *Em* β-catenin co-localizes with F-actin plaques that form where F-actin stress fibers are aligned between neighboring cells. This is consistent with the view that the AEP is a contractile epithelium (Elliott and Leys 2007; Nickel et al. 2011) and that the plaques represent AJs that form at focal points of tissue stress.

### A possible cell-substrate adhesion role for *Em* β-catenin

Detected interactions of *Em* β-catenin with *Em* α-catenin and *Em* CDH2, and its presence at cell-cell contacts, support predictions about its possible role in AJ in sponge tissues. A less expected result is the detection of *Em* β-catenin at structures in the attachment epithelium that are likely cell-substrate adhesions (Fig 4). These structures resemble focal adhesions (FAs) except that FAs utilize integrins rather than cadherins as adhesion receptors to mediate interactions between cells and the ECM. We found no evidence that FA components co-precipitate with *Em* β-catenin, and in bilaterians β-catenin is not a typical FA component. However, Langhe and colleagues (2016) have shown that *Xenopus* β-catenin and cadherin 11 co-localize with FA components when overexpressed in HeLa cells and Kuo and colleagues (2011) found β-catenin associated with FAs isolated from HFF1 cells. The functional significance of these studies is unknown, and thus the implications for our finding of *Em* β-catenin at cell-substrate adhesions in *E. muelleri* are unclear.

It is possible that classical cadherins mediate both cell-cell and cell-matrix adhesion in *E. muelleri*. Like classical cadherins in *Drosophila* and other non-chordate animals, *Em* CDH1 and *Em* CDH2 have membrane-proximal epithelial growth factor (EGF) and Laminin G (LamG) domains (Supplement Fig S2). In *Drosophila* this region has been implicated in trafficking to the plasma membrane (Oda and Tsukita 1999), but in an unusual classical cadherin-like protein (BbC) in the lancelet *Branchiostoma*, this region is sufficient to mediate epithelial cell adhesion [BbC naturally lacks cadherin repeats; (Oda et al. 2002; Oda et al. 2004)]. In general, EGF and LamG are involved in extracellular protein interactions and are common in both cell surface receptors and secreted components of the ECM. In the future, it will be critical to determine the expression dynamics and subcellular localization of candidate sponge adhesion receptors, themselves; both integrins and classical cadherins.

### Evidence for a functional Wnt/β-catenin pathway in sponges

Activation of the Wnt/β-catenin signaling pathway results in the stabilization of cytosolic β-catenin and its subsequent translocation to the nucleus (MacDonald et al. 2009; Cadigan and Waterman 2012). In *E. muelleri*, we find no evidence for conserved interactions of *Em* β-catenin with Wnt pathway components (such as TCF/Lef, Axin or APC) by Co-Immunoprecipitation, but by immunostaining we detect a nuclear population of *Em* β-catenin in cells of the attachment epithelium and in archeocytes (migratory cells within of the mesohyl) where it presumably functions as a transcriptional coactivator. Additionally, we find evidence that *Em* β-catenin is a substrate of GSK3B in *E. muelleri*, as inhibition of GSK3B leads to a shift in endogenous *Em* β-catenin phosphorylation states. This resembles the condition in bilaterians, where GSK3B functions as a negative regulator of Wnt signaling by phosphorylating free cytosolic β-catenin, leading to its ubiquitination and degradation via the proteasome (MacDonald et al. 2009; Stamos and Weis 2013).

Archeocytes are hypothesized to be pluripotent cells. Part of the evidence for this perspective is that they express stem cell markers such as *Musashi* and *Piwi* (Alié et al. 2015), and disruption of archeocyte cell division inhibits differentiation of the choanoderm (Rozenfeld and Rasmont 1977; Peña et al. 2016). Thus, detection of nuclear *Em* β-catenin in archeocytes (Fig 5; migratory cells in mesohyl are mostly archeocytes) is consistent with the known role of Wnt/β-catenin signaling in bilaterian stem-cell maintenance and renewal. Wnt/β-catenin pathway genes expression has also been reported in archeocytes of the sponges *A. queenslandica* (Adamska et al. 2010) and *S. ciliatum* (Leininger et al. 2014). If β-catenin has a conserved role in maintaining archeocyte pluripotency in *E. muelleri*, it may explain aspects of the aquiferous system phenotypes observed with GSK3B inhibition. Specifically, sustained activation of the Wnt/β-catenin pathway may inhibit differentiation of archeocytes into choanocytes.

A nuclear population of *Em* β-catenin is also present in cells of the attachment epithelium. One possible explanation for this finding comes from a study implicating Wnt/β-catenin signaling as an upstream regulator of collective cell migration in the migrating zebrafish lateral line primordium, particularly in cells of the leading edge of migration (Aman and Piotrowski 2008). It is well established that sponge are capable of entire-colony mobility, and in juveniles of *E. muelleri* this behavior is the result of collective migration of the attachment epithelium (Bond and Harris 1988; Bond 1992).

A limitation of our study is that we were restricted to working with gemmule-hatched juvenile tissues, whereas Wnt/β-catenin has previously been implicated in embryonic development in other sponge species. Specifically, in *A. queenslandica* (Adamska et al. 2010) and *S. ciliatum* (Leininger et al. 2014), Wnt/β-catenin pathway genes were found to exhibit posterior/anterior expression gradients during embryogenesis, consistent with a conserved role in axial patterning. These data are difficult to relate to our findings in *E. muelleri* because embryogenesis is a fundamentally different process than gemmule hatching, and the regulatory similarities between these disparate developmental processes have never been examined. Nonetheless, (Windsor and Leys 2010) also studied gemmule-hatched *E. muelleri* juveniles and reported that GSK3B inhibition disrupted development of the aquiferous system and caused the formation of ectopic oscula. From these data, they hypothesized that the adult body axis in sponges may be defined by the unidirectional aquiferous system. Whereas our data do not bear on this question directly, they do support the underlying assumption that *Em* β-catenin is a substrate of GSK3B in *E. muelleri*, and confirm that the optimal effective dosages of GSK3B inhibition of β-catenin phosphorylation recapitulate the reported knockdown phenotypes.

Finally, a gene expression study in *S. ciliatum* has also implicated Wnt/β-catenin signaling in endomesoderm specification, and revealed β-catenin expression in the embryonic micromeres (precursors of choanocytes) as well as mature choanocytes and mesohyl cells (Leininger et al. 2014). Based upon this study, it was hypothesized that sponges have germ layers homologous to those of bilaterians and that the choanoderm derives from endoderm. However, contradictory data from lineage-tracing experiments in *A. queenslandica* indicate that cell layers in sponges do not undergo progressive fate determination and are therefore not homologous to bilaterian germ layers (Nakanishi et al. 2014). Our data bear on this question only in that we detect a cell-adhesion-associated population of *Em* β-catenin in the choanoderm, but not a nuclear population. It is therefore possible that the β-catenin expression detected in the choanoderm of *S. ciliatum* relates to its roles in cell adhesion in this tissue rather than Wnt signaling. However, it is notable that the modern sponge lineages to which *E. muelleri* and *S. ciliatum* belong, diverged ~750-850 Ma, during the Neoproterozoic (Dohrmann and Wörheide 2017), and they have quite divergent developmental features.

## SUMMARY AND CONCLUSIONS

Here, we study β-catenin in the freshwater sponge *E. muelleri* and find 1) that AJ components (α-catenin and classical cadherin), but no Wnt pathway components, co-precipitate with endogenous *Em* β-catenin from whole-cell lysates, 2) there are cell boundary (adhesion-related) and nuclear (signaling-related) populations of *Em* β-catenin, 3) that previously reported phenotypic effects of GSK3B inhibitors on aquiferous system development can be replicated and are associated with shifts in *Em* β-catenin phosphorylation state, and 4) there is an *Em* β-catenin population associated with probable cell-substrate adhesion structures in the attachment epithelium, a function unknown in other animals.

The evidence presented here supports the underlying assumptions of previous studies: that β-catenin has conserved functions in Wnt signaling and cadherin-based cell adhesion in sponges. However, Wnt pathway components did not co-precipitate with *Em* β-catenin, and there is emerging evidence that Wnt/β-catenin is absent in hexactinellid sponges, warranting caution in interpretation (Schenkelaars et al. 2017). Furthermore, the discovery of a β-catenin population associated with cell-substrate adhesion structures underscores that functional conservation should not be uncritically accepted based upon bioinformatics and gene expression studies alone. Particularly, in the case of pleiotropic genes, the interpretation of gene expression patterns is potentially fraught without more explicit functional and biochemical support. To this end, future progress will depend largely upon the development of experimental approaches to manipulate gene function directly, *in vivo*, without the ambiguities that accompany pharmacological approaches. Moreover, comparative research should continue in multiple sponge species, as modern sponges represent many anciently divergent lineages.

## MATERIAL AND METHODS

### Collection and cultivation of sponges

Adult specimens of *E. muelleri* containing gemmules were collected in October 2013 from upper Red Rock Lake (Brainard Lake Recreation Area), Colorado, USA. The gemmule-bearing sponges were stored in autoclaved lake water in the dark at 4 °C and water was periodically refreshed. Gemmules were removed from the adult by gentle rubbing of the sponge tissue or by picking gemmules from the tissue with forceps. To clean the gemmules, they were washed several times with autoclaved lakewater.

For obtaining sponge cell lysate for immunoprecipitation, ~100 sponges were cultivated in a 90 mm petridish containing 25 ml of autoclaved lake water or sterile M-medium [1 mM CaCl_2_·6H_2_O, 0.5 mM MgSO_4_·7H_2_O, 0.5 mM NaHCO_3_, 0.05 mM KCl, 0.25 mM Na_2_SiO_3_; (Funayama, et al. 2005)] in the dark at room temperature (RT). Sponges typically hatched 4-5 days after plating, and lakewater or M-medium was refreshed every other day. Three to four days after hatching the sponges were fully developed and used to prepare cell lysates.

To culture sponges for immunostaining and pharmacological studies, gemmules were placed on glass coverslips (#1.5, 22 mm x 22 mm) in a 6-well plate containing 5 mL of autoclaved lakewater or M-medium per well. Sponges were incubated for 3-4 days at RT in the dark, and after hatching were fixed for immunostaining or analyzed by Western Blot.

### cDNA library construction

Material for constructing the cDNA library was obtained from juvenile sponges derived from one adult specimen of *E. muelleri*. Total RNA (~1 mg) was isolated using TRIzol Reagent (ThermoFischer Scientific #15596026) following manufacturer’s instructions. mRNA was purified from total RNA by poly A selection with the Oligotex Suspension kit (Qiagen #79000) and concentrated using Corning Spin-X UF 500 µl columns with a molecular weight cut-off of 10,000 (Corning #431478). To construct the cDNA library for use as a PCR template in cloning, the CloneMiner II cDNA Library Construction Kit was used (ThermoFischer Scientific #A11180) according to the manufacturer’s instructions.

### Protein expression and purification

The gene region encoding the N-terminus of *E. muelleri* β-catenin (*Em* β-catN amino acids 1-221) was amplified by PCR from *E. muelleri* cDNA with the following primers:

*Em* β-catN forward:

1) tacttccaatccaatgcaATGGAGGTGGACAGATCATACTAC, which contains an LIC adapter (lower case) *Em* β-catN reverse
2) ttatccacttccaatgttatta<u>CTA</u>GAGCTCGGAGGAACCCTGAT, which contains an LIC adapter (lower case) and a STOP codon (underlined).

The amplified fragment was cloned into the pET His6 Sumo TEV LIC cloning vector (1S) (Addgene plasmid # 29659). This vector generates proteins that are N-terminally tagged with a TEV-cleavable His6-Sumo fusion tag. The construct was transformed into *E. coli* Rosetta (DE3) competent cells (Novagen). Recombinant protein expression was induced at OD600 ~0.4 by the addition of 0.25 mM IPTG and was conducted at room temperature for 6 hours. Cells were spun down and resuspended in cell lysis buffer (1x PBS, 2M NaCl, 5% glycerol, 0.1 mM PMSF, pH 8.0) containing lysozyme (1mg/ml) and incubated at room temperature for 10 min. The lysis solution was then sonicated by 4 x 30sec pulses and insoluble bacterial debris was removed by centrifugation at 20,000xg for 30 min at 4 °C. To purify the recombinant protein, the soluble fraction of the *E. coli* lysate was incubated with HisPur Cobalt Resin (ThermoFisher # 89964) overnight at 4 °C. The resin was washed with washing buffer (1xPBS, 5% glycerol, pH 8.0) and the His-tagged proteins were eluted with elution buffer (150 mM imidazole, 1 x PBS, 5% glycerol, pH 8.0). To cleave the His6-Sumo fusion tag, recombinant proteins were incubated with AcTEV protease (ThermoFischer # 12575015) overnight at 4 °C, using 1 U for ~40 µg of recombinant protein. To remove imidazole and DTT, a buffer exchange was performed using PD-10 Desalting columns (GE Healthcare # 17-0851-01) and the protein samples were equilibrated in 1x PBS (pH 8.0) with 5% glycerol. Subsequently HisPur Cobalt resin was used to remove the His6-Sumo tag and AcTEV protease. The eluate contained the *Em* β-catN recombinant protein (without His6-Sumo fusion tag), which was used to inject in rabbits.

### Antibody Production

Polyclonal antibodies were raised in rabbits against recombinant *Em* β-catN antigen (Syd labs, Natick, USA) and the anti-Em β-catN antibodies were affinity purified from the antiserum using recombinant *Em* β-catN protein with the AminoLink Plus Immobilization Kit (ThermoFischer, #44894) according to the manufacturer’s instructions. Two Amino-link columns were prepared, one with *Em* β-catN recombinant protein and one with *E. coli* lysate. First the rabbit serum was incubated with the *E.coli* lysate-column by running it 10x over the column to remove possible IgGs against bacterial proteins. Next, the eluate was incubated with the *Em* β-catN protein-column by running it 10x over the column and the column was washed extensively with 1x PBS. Finally, the affinity-purified antibody was eluted from the column with 0.1 M Glycine (pH 2.5) and neutralized with 0.75M Tris (pH 8.8). The fractions with the highest antibody concentrations were combined, and buffer was exchanged with PD-10 Desalting columns and the affinity purified antibody was equilibrated in 1x PBS containing 0.05% sodium azide. The final concentration of affinity purified antibody was ~1 mg/ml.

### Immunoprecipitation and Mass Spectrometry

Two independent *Em* β-catenin immunoprecipitation experiments were performed using Dynabeads Protein G for Immunoprecipitation (ThermoFischer Scientific, # 10004D) according to the manufacturer’s instructions. For each experiment, 5 µg of *Em* β-catN-antibody was coupled to 50 µl (=1.5 mg) of Dynabeads protein G. Normal rabbit IgG (Santa Cruz Biotechnology, #sc-2027) was coupled to Dynabeads as a negative control. To crosslink the antibody to the beads, the antibody-coupled beads were incubated with 350 µl of 0.75 mM BS3 crosslinker (ThermoFischer Scientific, #21580) in conjugation buffer (20 mM Sodium Phosphate, 0.15M NaCl, pH 8.1) for 30 min at room temperature, crosslinking reaction was stopped by adding 17.5 µl Quenching buffer (1M Tris-HCl, pH 7.5), and the beads were washed with PBST (0.1% Tween-20).

Whole cell lysates were derived from ~1 week old gemmule-hatched juveniles. At this stage, all somatic tissues had developed. The lysate was prepared by dissolving the juvenile sponges in lysis buffer (20 mM Hepes, pH 7.6, 150 mM NaCl, 1 mM EDTA, 10% Glycerol, 1% Triton X-100, 1 mM DTT, 1 mM PMSF, 1x phosphatase inhibitor cocktail A and B (Biotool, #B15001), 1x protease inhibitor cocktail (Biotool, #B14011, EDTA free)). The crude lysate was clarified by centrifugation at 14,000xg for 10min at 4 °C. From the crude lysate we took 600 µg of soluble sponge proteins (measured with Bradford Protein Assay) and diluted this 2x with binding/washing buffer (same as lysis buffer, but without glycerol and DTT). The lysate was incubated with the antibody-coupled beads for 1.5 hr at 4 °C, and was washed with binding/washing buffer and proteins were eluted with 50 µl of 0.2 M Glycine pH 2.5.

The samples were digested, desalted and analyzed with mass spectrometry by EPFL-Plateforme technologique de proteomique using Orbitrap Elite (4x short LC-MS/MS gradient). The raw data was searched against the *E. muelleri* protein database using Scaffold software with a protein threshold of 99% and a peptide threshold of 95%. These results were compared with the control-immunoprecipitation (rabbit IgG) to identify non-specific interactions of sponge proteins with IgG-coupled Dynabeads. Proteins that had at least 5 unique peptide matches and no hits in the control-immunoprecipitation were selected (see Table 1. For a complete list of all identified proteins, see Supplement Table S2).

### Immunostaining

Sponges that grew on the coverslips in 6-wells format plates were fixed with ice-cold EtOH with 4% formaldehyde for 1h at 4 °C. To remove the fixative, the sponges were washed 3x with PBST (0.1% Tween-20). The coverslips containing the sponges were transferred to parafilm in a humid chamber. First sponges were incubated with blocking reagent (3% BSA in PBST (0.1% Tween-20)) for 1 h at room temperature. Next, they were incubated with the primary antibody (anti-*Em* β-catN 1:300 in blocking reagent) for 2 hrs at room temperature or overnight at 4 °C. After incubating with the primary antibody, sponges were washed 3x with PBST and were incubated with the secondary antibody (1:500, goat anti-rabbit Alexa Fluor 488, ThermoFischer #A11034) together with Alexa Fluor 568 Phalloidin (1:40, ThermoFischer #A12380) and Hoechst33342 (1 µg/ml) in blocking reagent for 45 min at room temperature. Sponges were washed 3x with PBS before they were mounted with mounting medium (90% glycerol, 1x PBS, 0.1 M propyl gallate) on a microscopic glass slide with clay feet and sealed into place with heated VALAP (vaseline, lanolin and paraffin, 1:1:1). The negative controls were incubated with secondary antibody only. Images were taken with a 60x and 100x oil immersion objective on an Olympus Fluoview 1000 inverted confocal microscope, and images were processed using ImageJ (z-stack: maximal intensity projection).

### Pharmacological treatment (AP and BIO) and Western blot analysis

All treatments were done in sterile M-medium with different concentrations of AP (Alsterpaullone, Sigma-Aldrich #A4847) and BIO (6-bromoindirubin-3′-oxime, Sigma-Aldrich #B1686). The tested concentrations for AP were 0.025 µM, 0.05 µM, 0.1 µM and 0.2 µM and for BIO 0.125 µM, 0.25 µM, 0.5 µM and 0.1 µM. Since AP and BIO were initially dissolved in DMSO (10 mM stocks), the control treatments consisted of 0.02% DMSO in M-medium. Sponges were treated for the duration of the experiment and were imaged during day 5, 6 and 7 using a Widefield Zeiss Cellobserver HS. At day 7, sponges were harvested for Western Blot analysis. To prepare samples for western blot, sponges were dissolved in 1x Lysis buffer and samples were boiled together with Laemmli loading buffer for 5 min, followed by a quick spin. The protein samples (~2 sponges per lane) were separated by SDS-PAGE on an 8% gel and the proteins were transferred to a PVDF membrane (Bio-Rad #1620177). Membranes were blocked for 1 h at room temperature in blocking solution (5% non-fat dry milk in 1x PBST (0.1% Tween-20)), and then incubated with anti-*Em* β-catN antibody (1:10.000) in blocking solution overnight at 4 °C Membranes were washed extensively with PBST and incubated with the secondary antibody (goat anti-rabbit IgG antibody, horseradish peroxidase conjugate, Promega #W401B 1:5000) in blocking solution for 1h at room temperature. Again, membranes were washed extensively in PBST and were developed using Western Lightning Plus Enhanced Chemiluminescence Substrate (PerkinElmer #NEL104001EA). To visualize the loading control, the membrane was stripped and re-probed. Stripping was done by incubating the membrane for 10 min at room temperature in mild stripping buffer (1.5% w/v glycine, 0.1% w/v SDS and 1% w/v Tween-20) and washing extensively in PBS and PBST. Re-probing procedure was similar as described above, but using mouse anti-alpha tubulin (1:10.000, Sigma Aldrich #T5168) as a primary antibody and goat anti-mouse rabbit IgG antibody, horseradish peroxidase conjugate (1:5000 Promega #W402B) as a secondary antibody.

## ACKNOWLEDGEMENTS

The authors thank Brigitte Galliot and Detlev Arendt for providing lab space, reagents and administrative support for KJS for periods of the study, and Jennyfer Mora Mitchell for help with field collection of *E. muelleri*. This work was supported by a Marie-Curie Fellowship (grant number FP7-PEOPLE-2012-IOF-327684 Sponge Signaling) [KJS] and the University of Denver Faculty Research Fund [SAN].

